# Mitochondrial functional resilience after TFAM ablation in adult cardiomyocytes

**DOI:** 10.1101/2020.06.18.159863

**Authors:** Nasab Ghazal, Jessica N. Peoples, Tahmina Mohuiddin, Jennifer Q. Kwong

## Abstract

The adult heart is a terminally differentiated tissue that depends on mitochondria for its energy supply. Respiratory chain energy supply deficits due to alterations in the mitochondrial genome (mtDNA) or in nuclear genome (nDNA)-encoded mtDNA regulators are associated with cardiac pathologies ranging from primary mitochondrial cardiomyopathies to heart failure. Mitochondrial transcription factor A (TFAM) is an nDNA-encoded regulator of mtDNA transcription, replication, and maintenance. Insufficiency of this protein in embryonic and postnatal cardiomyocytes causes cardiomyopathy and/or lethality, establishing TFAM as indispensable to the developing heart; its role in adult tissue has been inferred from these findings. Here, we provide evidence that challenges this long-standing paradigm using *Tfam* ablation in the adult heart. Unexpectedly, loss of *Tfam* in adult cardiomyocytes resulted in a prolonged period of functional resilience characterized by preserved mtDNA content, mitochondrial function, and cardiac function despite mitochondrial structural alterations and decreased transcript abundance. Remarkably, TFAM protein levels did not directly dictate mtDNA content in the adult heart, and mitochondrial translation was preserved with acute TFAM inactivation, suggesting a mechanism whereby respiratory chain assembly and function can be sustained, which we term ‘functional resilience’. Finally, long-term *Tfam* inactivation induced a coordinated downregulation of the core mtDNA transcription and replication machinery that ultimately resulted in mitochondrial dysfunction and cardiomyopathy. Taken together, adult-onset cardiomyocyte-specific *Tfam* inactivation reveals a striking resilience of the adult heart to acute insults to mtDNA regulatory mechanisms and provides insight into critical differences between the developing versus differentiated heart.

## INTRODUCTION

Mitochondria are critical to fuel cardiac contraction. These organelles occupy ~30% of the cardiomyocyte by volume and provide ~90% of the ATP that is utilized on a beat-to-beat basis (1, 2). Defects in mitochondrial energy production underlie many diseases that affect the heart, ranging from primary mitochondrial cardiomyopathies due to defects in the oxidative phosphorylation (OXPHOS) system, to heart failure and aging (3–6). Dissecting mechanisms regulating mitochondrial energy production systems therefore affords critical knowledge to understand basic cardiac physiology.

Mitochondria are under the dual control of the nuclear genome (nDNA) and the mitochondrial genome (mtDNA). While most of the ~1100 proteins of the mammalian mitochondrial proteome are specified by nDNA (7), mtDNA specifies 13 essential subunits of the respiratory chain, alongside two ribosomal RNAs (rRNAs) and 22 transfer RNAs (tRNAs) that are required for mtDNA protein translation (8). The mtDNA is distinct from and regulated independently of the nDNA, but shares similar organization into discrete DNA-protein structures, called nucleoids (9).

MtDNA nucleoids encompass proteins involved in mtDNA maintenance, replication, transcription, and translation (10–12). A key protein within the mtDNA nucleoid structure is the mitochondrial transcription factor A (TFAM), an nDNA-encoded mtDNA-binding protein with established roles in mtDNA packaging (13–15), transcription (16, 17), replication (18), and copy number maintenance (19, 20). In vivo studies confirm these critical roles of TFAM: systemic inactivation of murine TFAM causes early embryonic lethality [by embryonic day (E) 10.5] that is accompanied by mtDNA depletion and reduced mitochondria-encoded respiratory chain complex activity (21).

The essential nature of TFAM in vivo is mirrored in the developing embryonic and postnatal heart. Inactivation of *Tfam* in the mouse embryonic heart at E7.5 using the *Nkx2.5-Cre* recombinase results in lethality by E15.5, with the embryonic hearts marked by myocardial wall thinning (22). *Tfam* deletion in the skeletal muscle and heart mediated by the muscle creatinine kinase promoter-driven Cre (*Ckmm*-Cre), which initiates deletion at E13, triggers development of dilated cardiomyopathy with atrioventricular conduction blocks and death by 2 to 4 weeks postnatal age (23). This dilated cardiomyopathy phenotype and severely blunted survival can be recapitulated with cardiomyocyte-specific *Tfam* inactivation in the postnatal heart using Cre under the control of the alpha myosin heavy chain promoter (αMHC-Cre), which induces robust gene deletion in the postnatal ventricular cardiomyocytes (24, 25). Unifying these existing mouse models, TFAM ablation early in life causes severe mtDNA copy number reduction, impaired mtDNA gene expression, reduced mitochondrial respiratory chain complex assembly, respiratory chain dysfunction, and abnormal mitochondrial ultrastructure (22–25).

These existing models define the importance of proper mtDNA regulation in the developing heart, which is a period of active cardiomyocyte growth and proliferation (26). The role of mtDNA regulation in the adult heart has been inferred from such findings. However, the heart is a terminally differentiated tissue. The transition from the postnatal heart at birth to a fully developed adult organ is marked by multiple rounds of early cardiomyocyte proliferation followed by hypertrophic growth (26). During this period, cardiac mitochondria also undergo a profound structural and functional maturation, with the fragmented mitochondria of embryonic and early postnatal cardiomyocytes developing into the interconnected network of adult cardiomyocytes (27). This transition is marked by increase mitochondrial biogenesis and a metabolic shift toward β-oxidation (27, 28). The fully differentiated adult heart could therefore have different needs for mtDNA maintenance, but this question has not been investigated.

In this study, we investigated the function of the mtDNA maintenance machinery in the adult heart by engineering a mouse model with tamoxifen-inducible cardiomyocyte-specific deletion of *Tfam* (*Tfam^fl/flxMCM^* mice). Surprisingly, we found that TFAM does not determine mtDNA copy number in the adult heart and that the adult heart displays a remarkable functional resilience in response to TFAM ablation, where cardiac structure, function, and mitochondrial energetics can be maintained. Moreover, mitochondrial function appears to be sustained by preserved mitochondrial protein translation. Finally, while long-term (>5 months) TFAM ablation causes dilated cardiomyopathy marked by mtDNA depletion as well as reduced mtDNA gene expression, these phenotypes may be potentiated by a coordinated downregulation of the mitochondrial transcription and replication machinery. Collectively, our data demonstrate differing responses to defects in the mtDNA maintenance and expression machinery between developing/postnatal and adult hearts, and point to preserved mitochondrial translation as a potential pathway that confers resistance to mitochondrial insults in the adult myocardium.

## RESULTS

### Cardiomyocyte-restricted ablation of TFAM in the developing early postnatal versus adult mouse heart

To dissect effects of mtDNA machinery disruptions in the mouse heart, we compared two models of cardiomyocyte-specific ablation of TFAM. For postnatal deletion of TFAM in the heart, we crossed *Tfam* loxP-targeted animals (*Tfam^fl/fl^*) (21) with mice expressing a Cre recombinase under the control of the cardiomyocyte-specific αMHC promoter (Cre; *Tfam^fl/flxCre^*, Figure 1A) (29). To achieve deletion of TFAM in the adult heart, we crossed *Tfam^fl/fl^* mice with mice harboring a tamoxifen-inducible αMHC-driven Cre (MCM; *Tfam^fl/flxMCM^*, Figure 1E) (30). Cardiomyocyte-specific TFAM deletion was induced by systemic tamoxifen administration to adult (8-week-old) *Tfam^fl/flxMCM^* experimental mice, with *Tfam^fl/fl^* mice serving as controls (tamoxifen 25 mg/kg/day for 5 days, Figure 1F). Both the constitutive postnatal αMHC-Cre (evaluated at 15 days of age) as well as the inducible adult MCM systems (evaluated 2 weeks post-tamoxifen administration, Figure 1B) resulted in a significant reduction in TFAM protein expression in the heart as assessed by immunoblotting (Figure 1G). TFAM protein levels were not further reduced in *Tfam^fl/flxMCM^* versus *Tfam^fl/fl^* cardiac tissue by extending the time between tamoxifen dosing and immunoblot analysis of protein expression (data not shown), suggesting that 2 weeks following the tamoxifen-initiated Cre recombination maximally decreases TFAM in the heart.

**Figure 1.**
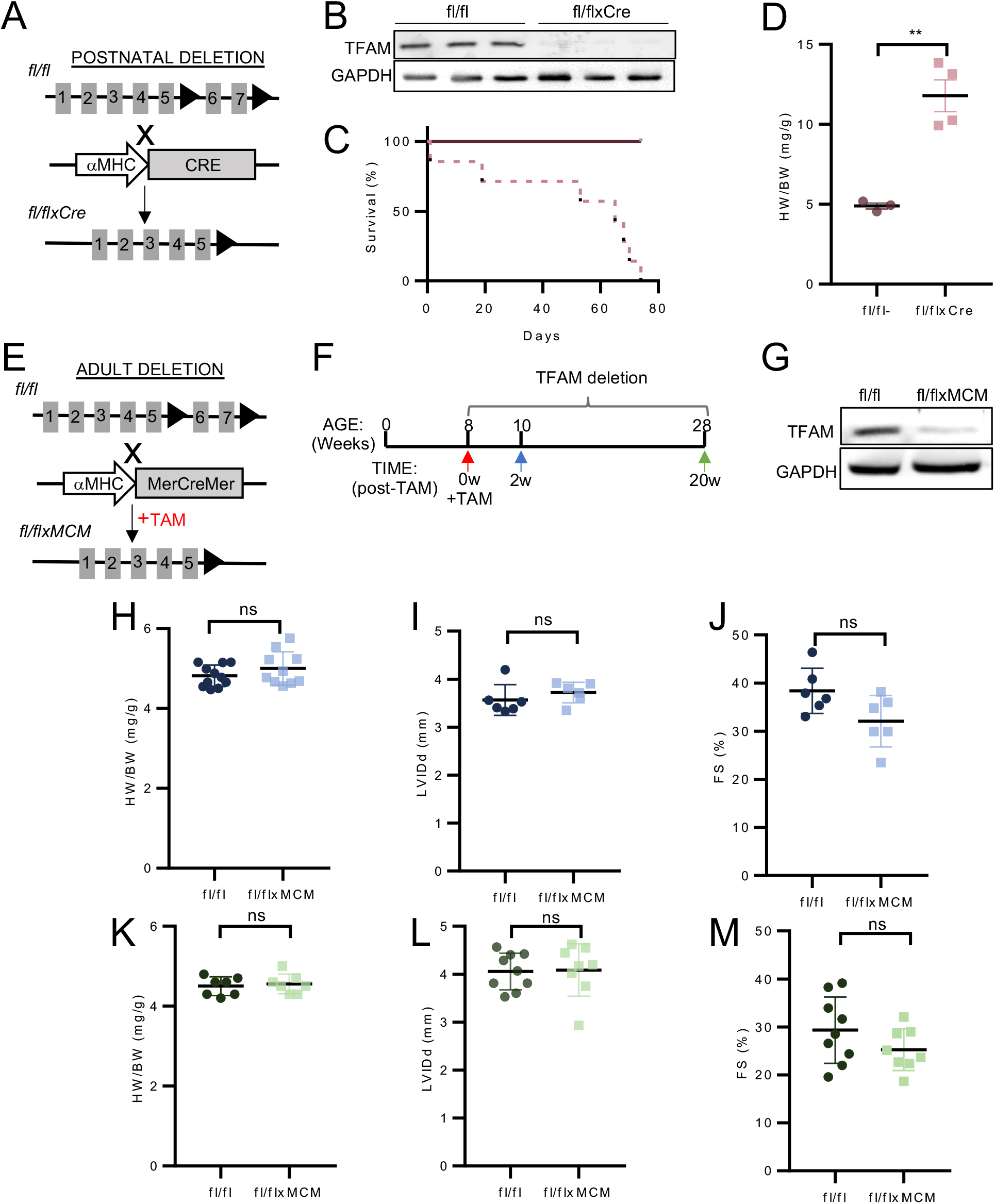
Acute loss of TFAM has no structural or functional impact on the adult mouse heart. (A) Generation of the *Tfam^fl/flxCre^* model of TFAM deletion in postnatal cardiomyocytes. Tfam loxP (red triangles) targeted mice (fl/fl) crossed to mice expressing an αMHC promoter-driven Cre recombinase. B) Western blot of TFAM expression in total cardiac protein of the indicated genotypes at 15 days of age. GAPDH was used as a loading control. (C) Kaplan-Meier survival curve (n=7/group). (D) Heart weight to body weight ratio (HW/BW) of the indicated groups of mice at 60 days of age. **P<0.01 versus fl/fl mice. (E) Generation of the *Tfam^fl/flxMCM^* mice. *Tfam^fl/fl^* mice were crossed to αMHC-MerCreMer (MCM) transgenic mice, which express a tamoxifen (TAM) inducible Cre under the control of the αMHC promoter. (F) Tamoxifen dosing regimen (25 mg/kg/day, *i.p*. for 5 days) and experimental time course for acute TFAM deletion (2- and 20-weeks post-tamoxifen). (G) Western blot of TFAM expression in adult cardiomyocytes isolated from *Tfam^fl/fl^* and *Tfam^fl/flxMCM^* animals at the 2 weeks post-tamoxifen time point. HW/BW, left ventricular interior dimension in diastole (LVIDd) and fractional shortening (FS) as assessed by echocardiography at 2 weeks (H-J) and 20 weeks (K-M) post-tamoxifen administration. All values reported as mean ± SEM. Student’s t test was used for statistical analysis. P<0.05 were considered significant.

### Acute loss of TFAM has no structural or functional impact on the adult mouse heart

TFAM deletion systemically or in the embryonic heart causes embryonic lethality, while deletion in either developing striated muscle or cardiomyocytes results in mitochondrial cardiomyopathy and early lethality (22, 24, 25). In line with these previous studies, we found that *Tfam^fl/flxCre^* mice exhibited reduced survival, with no animals surviving longer than 74 days of age (Figure 1C), as well as a significant increase in heart weight/body weight ratios as compared to *Tfam^fl/fl^* controls at 60 days of age (Figure 1D). Thus, unsurprisingly, postnatal TFAM deletion in the heart induces pathological cardiac remodeling and hypertrophy.

To investigate the functional consequences of TFAM ablation in the adult heart, gravimetric and echocardiographic analyses were initially conducted on TFAM-deleted (*Tfam^fl/flxMCM^*) and control (*Tfam^fl/fl^*) mice 2 weeks after tamoxifen administration. At this acute time point, despite the significant decrease in TFAM protein levels in cardiomyocytes (Figure 1G), no changes were detected in gravimetric or echocardiographic parameters (Figures 1H–1J). Strikingly, cardiac dimensions and function (heart weight/body weight ratios, left ventricular interior dimensions, fractional shortening) remained unaltered even at 20 weeks after tamoxifen dosing (Figures 1K–1M). Together, these data suggest that, unlike in the postnatal developing heart, in the adult heart following acute loss, TFAM is dispensable for structural or functional maintenance of the heart.

### TFAM-deleted animals develop cardiomyopathy with aging

The strikingly normal cardiac structure and function following acute loss of TFAM in adult hearts prompted us to investigate the effects of prolonged TFAM loss. At 32 weeks after tamoxifen dosing (Figure 2A), histological analyses of heart sections revealed extensive inflammatory cell infiltrates (Figure 2B), significantly elevated cardiomyocyte cross-sectional area (Figure 2C), and increased fibrosis (Figure 2D) in *Tfam^fl/flxMCM^* mice as compared to *Tfam^fl/fl^* controls. Additionally, *Tfam^fl/flxMCM^* mice developed elevated heart weight/body weight and lung weight/body weight ratios as compared to *Tfam^fl/fl^* controls (Figures 2D and 2E), indicating cardiac hypertrophy and pulmonary edema. Echocardiographic analyses further showed that TFAM-deleted animals had increased left ventricular dimensions at diastole (LVIDd; Figure 2F) and decreased fractional shortening (FS; Figure 2G), indicating ventricular dilation and impaired cardiac performance. Collectively, these data show that *Tfam^fl/flxMCM^* mice develop dilated cardiomyopathy with aging.

**Figure 2.**
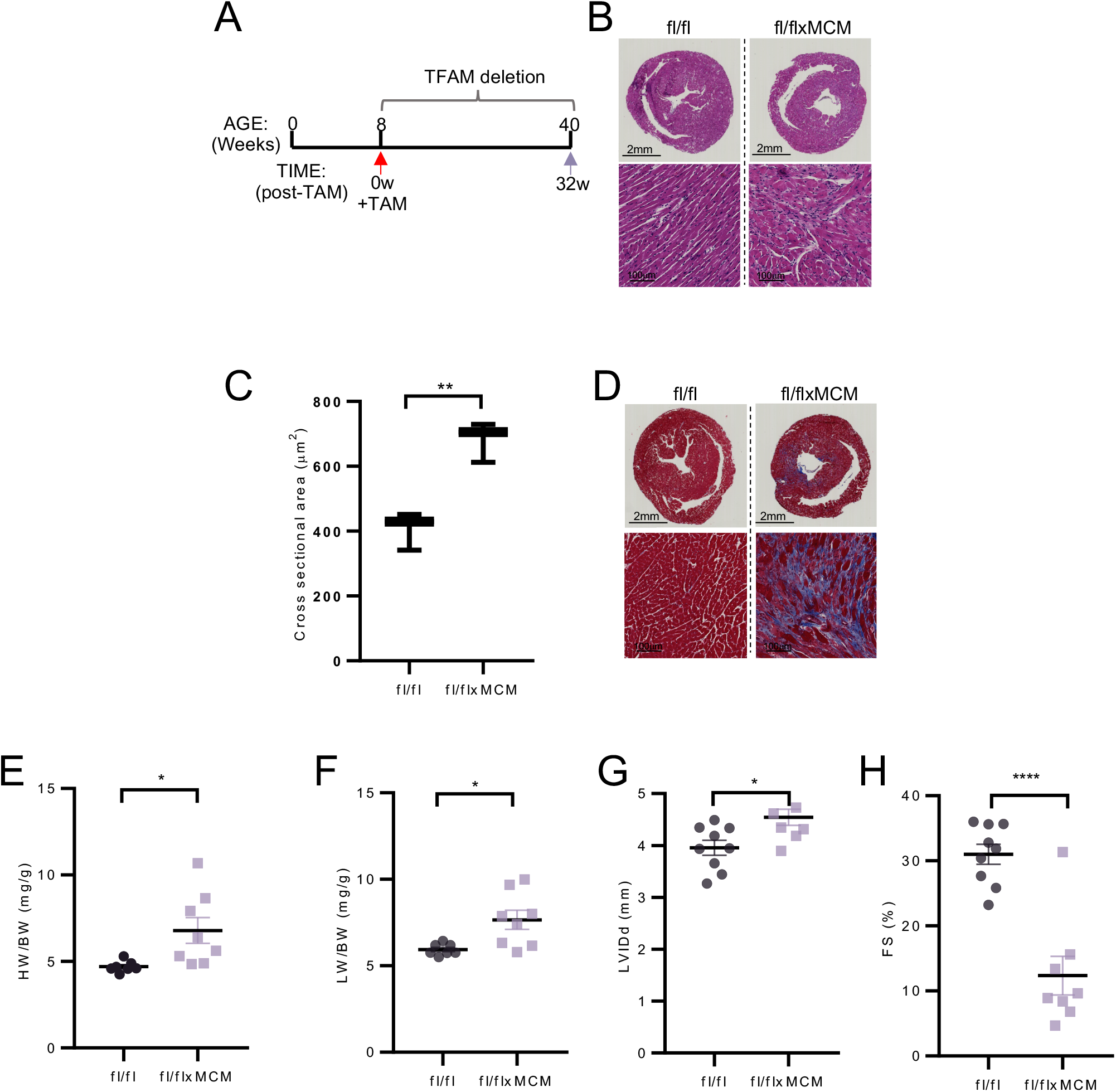
Prolonged TFAM inactivation in adult cardiomyocytes causes a delayed-onset mitochondrial cardiomyopathy. (A) Tamoxifen dosing and experimental time course for prolonged TFAM deletion (32 weeks post-tamoxifen). Representative images of 7-μm transverse heart sections from mice of the indicated genotypes at 32 weeks post-tamoxifen stained with (B) H&E. *Tfam^fl/flxMCM^* hearts display extensive inflammatory infiltrate. (C) Quantification of myocyte cross-sectional area from the indicated genotypes (n=3 animals/group; >200 myocytes/animal evaluated). (D) Representative images of Masson’s trichrome staining. *Tfam^fl/flxMCM^* hearts display increased fibrosis as compared to controls. (E) HW/BW ratio, (F) lung weight to body weight ratio (LW/BW), (G) LVIDd, and (H) fractional shortening (FS) assessed at the 32 weeks post-tamoxifen time point. All values reported as mean ± SEM. Student’s t test was used for statistical analysis. P<0.05 were considered significant. P<0.05 (*) and P<0.0001 (****).

### Mitochondrial structural remodeling is an early response to TFAM deletion

To understand how TFAM ablation affects mitochondria in the adult heart, we examined cardiac respiratory chain function. Respirometry was conducted on mitochondria isolated from *Tfam^fl/flxMCM^* and *Tfam^fl/fl^* hearts at the 2-, 20-, and 32-week post-tamoxifen time points (Figures 3A-C). Consistent with our observations of maintained cardiac structure and function upon acute TFAM deletion (Figures 2H-M), no significant differences were observed in basal or maximal oxygen consumption rates in TFAM-deleted versus control mitochondria at either the 2- or 20-week deletion time points (Figures 3A and 3B). In contrast, TFAM deletion-induced deficits in respiration were observed at 32 weeks of TFAM ablation (Figure 3C). Thus, mitochondrial function can be maintained in the acute absence of TFAM, but long-term TFAM deletion induces respiratory defects.

**Figure 3.**
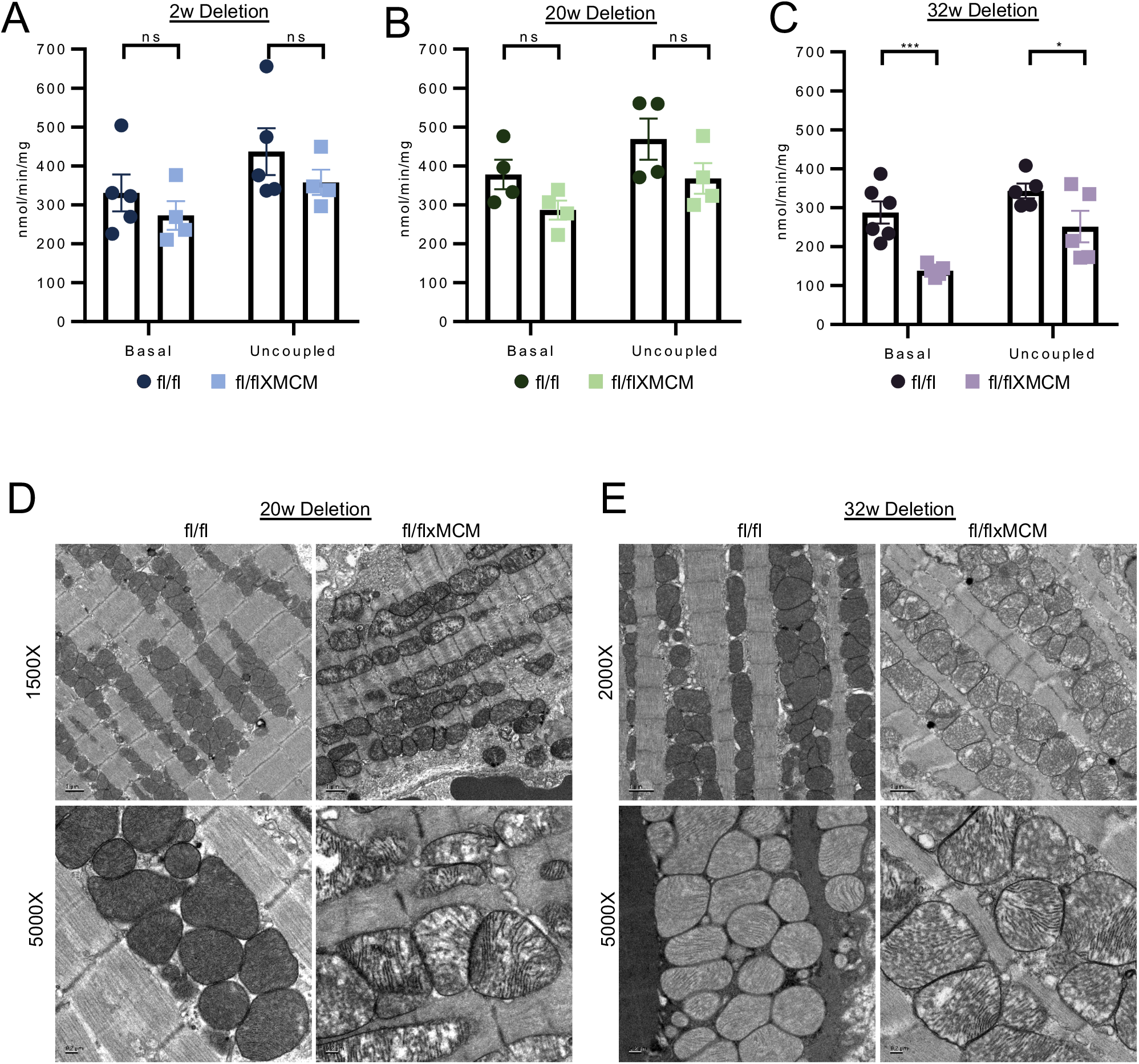
TFAM deletion-induced mitochondrial remodeling precedes onset of mitochondrial dysfunction. Basal and uncoupled mitochondrial oxygen consumption rates measured in isolated cardiac mitochondria from *Tfam^fl/fl^* versus *Tfam^fl/flxMCM^* mice at (A) 2 weeks, (B) 20 weeks and (C) 32 weeks post-tamoxifen. (D) Representative electron micrographs of left ventricular ultrastructure from the genotypes indicated at 20 weeks and (E) 32 weeks post-tamoxifen. All values presented as mean ± SEM. Student’s t test was used for statistical analysis. P<0.05 were considered significant. P<0.05 (*) and P<0.001 (***).

To determine the effects of TFAM loss on mitochondrial organization, transmission electron microscopy was conducted to examine mitochondrial ultrastructure. Remarkably, TFAM deletion disrupted cardiac mitochondrial morphology as early as 20 weeks after tamoxifen administration (Figures 3D and 3E). Importantly, TFAM deletion-induced alterations in mitochondrial organization (20 weeks; Figure 3D) occurred before the onset of respiratory chain dysfunction (observed at 32 weeks, Figure 3C). Together, these observations support a model whereby TFAM participates in the maintenance of cardiac mitochondrial organization, and this structural role of TFAM appears distinct from the effects of TFAM loss on mitochondrial energetics.

### TFAM deletion-induced mtDNA transcription defects precede mtDNA depletion

Deletion of TFAM in vitro, as well as in the contexts of embryonic and early postnatal cardiac development, causes respiratory chain dysfunction due to mtDNA depletion, impaired transcription, and a decline in respiratory chain complex expression and assembly (21, 23–25). Our observation that adult heart mitochondria display preserved respiration (Figures 3A and 3B) for months in the absence of TFAM protein was therefore unexpected. Given this surprising finding, we examined mtDNA content and mtDNA transcript levels in *Tfam^fl/flxMCM^* versus *Tfam^fl/fl^* hearts. Steady-state levels of representative mitochondrial transcripts spanning different mtDNA-encoded respiratory chain complexes (MTND5 and MTND6 for complex I, MTCYTB for complex III, MTCO1 for complex IV, and MTATP6 for complex V) were assessed by real-time PCR (RT-PCR). TFAM deletion in the heart resulted in a progressive decline in mtDNA-encoded transcripts, with decreased abundance of some transcripts *(Mtnd5)* observed as early as 2 weeks post-tamoxifen administration (Figures 4D-F). This alteration suggests that loss of TFAM induces an immediate decline in mtDNA transcription. Moreover, transcript expression of the mitochondrial RNA degradosome constituents *Pnpt1* (mitochondrial polyribonucleotide nucleotidyltransferase 1) and *Supv3l1* (mitochondrial ATP-dependent RNA helicase), which participate in mitochondrial RNA degradation (31), were unchanged at the 2- and 20-weeks post-tamoxifen times points (Supplemental Figure 1), suggesting that changes in mitochondrial RNA half-life do not contribute to TFAM deletion-induced decrease in mtDNA transcript levels.

**Figure 4.**
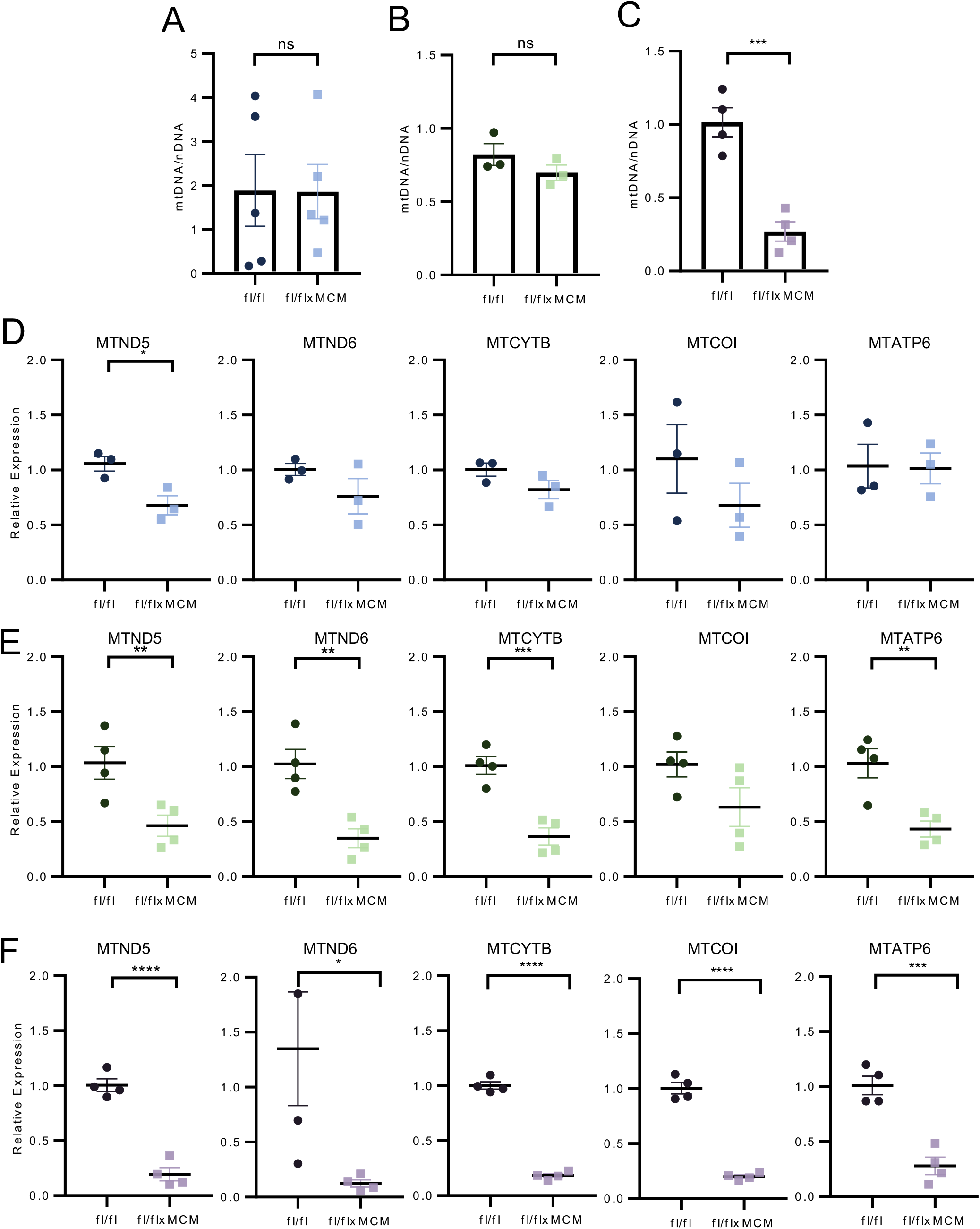
Defects in mtDNA transcription are an early response to TFAM deletion. Quantification of cardiac mtDNA to nDNA ratios from animals at the (A) 2-, (B) 20-, and (C) 32-week post-tamoxifen time points. RT-PCR analyses of steady state mtDNA-encoded transcript levels at (D) 2-, (E) 20-, and (F) 32-weeks post-tamoxifen. Values presented as mean ± SEM. Student’s t test was used for statistical analysis. P<0.05 were considered significant. P<0.05 (*), P<0.01 (**), P<0.001 (***), P<0.0001 (****).

To evaluate the impact of TFAM deletion on mtDNA content, mtDNA/nDNA ratios were measured from total DNA isolated from hearts of *Tfam^fl/flxMCM^* versus *Tfam^fl/fl^*. Surprisingly, decreased mtDNA/nDNA ratio was observed only in *Tfam^fl/flxMCM^* hearts 32 weeks following tamoxifen administration (Figures 4A-C), suggesting that mtDNA content can be maintained for months in the absence of TFAM. Collectively, these data suggest that in the adult heart, there is a differential requirement for TFAM’s functions in mtDNA maintenance versus transcription, and these functions can be differentiated; there may be an ongoing requirement for TFAM in transcription, but TFAM’s roles in mtDNA maintenance and replication may only be appreciable with long-term deletion.

### Mitochondrial translation is maintained during functional resilience

Given that the early phase of TFAM deletion (between 2- and 20-weeks post tamoxifen, a period which we have termed ‘functional resilience’) is marked by preserved mitochondrial function despite decreases in transcription, we next asked if mitochondrial protein translation could be maintained despite decreases in transcription. In organello translation assays were performed on cardiac mitochondria isolated from mice (*Tfam^fl/fl^* and *Tfam^fl/flxMCM^*) 13 weeks post-tamoxifen. Quantification of the levels of *Tfam^fl/flxMCM^* versus *Tfam^fl/fl^* mitochondrial protein synthesis products revealed no differences between the two groups (Figure 5A-B). To further examine the consequences of TFAM ablation on respiratory chain complex assembly during this acute/early deletion phase, we performed blue native gel electrophoresis of *Tfam^fl/fl^* and *Tfam^fl/flxMCM^* mitochondria. Consistent with our observations of unaltered translation, no differences in respiratory chain complex abundance were detected between the two groups (Figure 5C-D). Collectively, these data indicate that, despite TFAM deletion and reduced steady-state mtDNA transcripts, *Tfam^fl/flxMCM^* myocytes can maintain normal levels of mitochondrial protein synthesis. The findings suggest a possible mechanism whereby mitochondrial function can be preserved despite impaired mtDNA gene expression in the fully differentiated cardiomyocyte.

**Figure 5.**
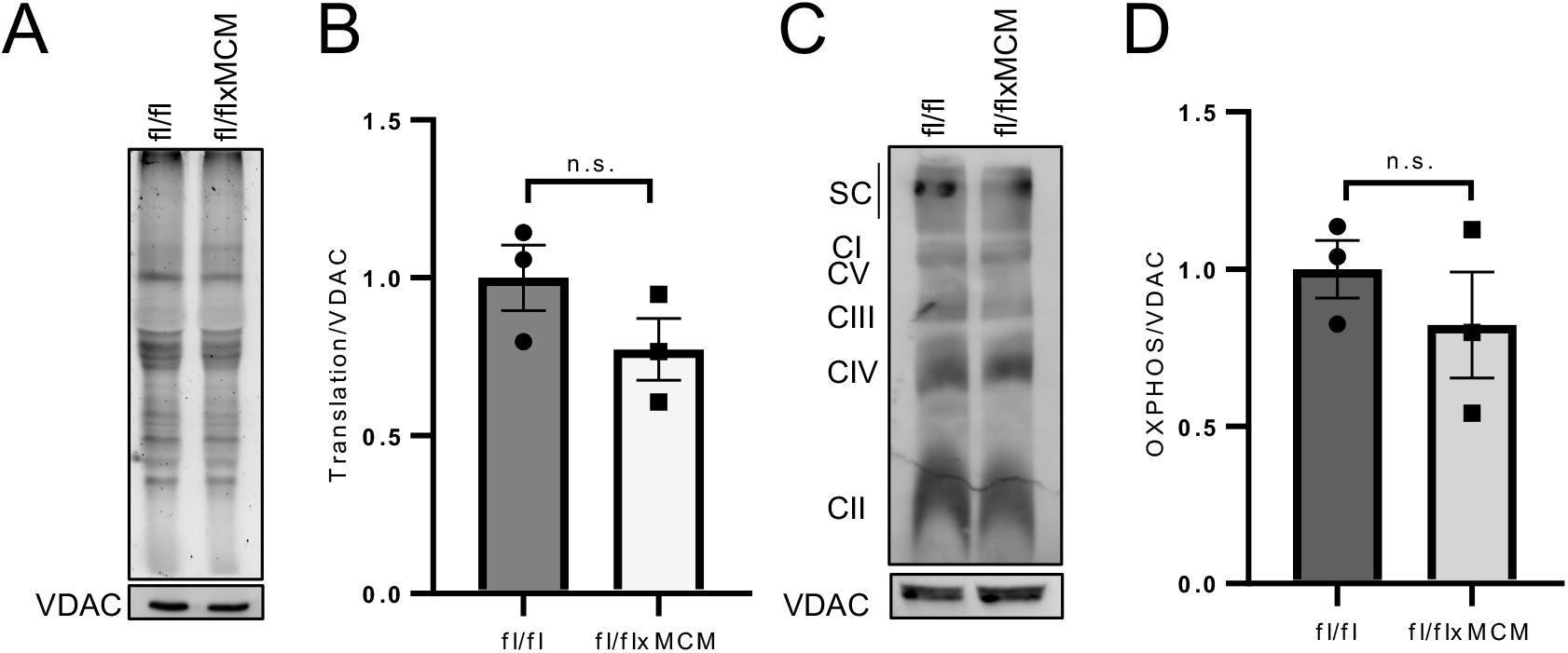
Mitochondrial protein translation is preserved during functional resilience. (A) Representative image of SDS-PAGE analyses of AHA-labelled proteins produced by in organello translation assays performed on cardiac mitochondria isolated from *Tfam^fl/fl^* and *Tfam^fl/flxMCM^* animals 13 weeks following tamoxifen dosing. A western blot for VDAC on the input mitochondria was used as a protein loading control. (B) Densitometry quantification of the in organello translation assays. (C) Analyses of the respiratory chain complex abundance and assembly in cardiac mitochondria isolated from *Tfam^fl/fl^* and *Tfam^fl/flxMCM^* animals 13 weeks post-tamoxifen. Mitochondrial proteins were resolved by BN-PAGE followed by immunoblotting with the Total OXPHOS antibody cocktail to simultaneously visualize all five complexes of the respiratory chain. A western blot for VDAC on the input mitochondria was used as a protein loading control. (D) Densitometry quantification of BN-PAGE. Values presented as mean ± SEM. Student’s t test was used for statistical analysis. P<0.05 (*).

### Prolonged TFAM loss causes a coordinated downregulation of mitochondrial transcription and replication machinery

Having established that translation is maintained in mitochondria in the absence of TFAM, possibly serving to maintain mitochondrial function (Figures 3A-B), we next investigated if additional factors ultimately contribute to the reduced mtDNA content (Figure 4C) and transcription (Figure 4F) as well as functional decline (Figure 3C) observed with prolonged TFAM ablation. Specifically, we focused on the mitochondrial transcription and replication machinery. Together with TFAM, mitochondrial transcription requires the coordinated actions of mitochondrial RNA polymerase (POLRMT), mitochondrial transcription factor B2 (TFB2M), and mitochondrial transcription elongation factor (TEFM) (32). Expression of these mtDNA transcription regulators was evaluated by RT-PCR. While *Polrmt, Tfb2m*, and *Tefm* transcript levels were unchanged in *Tfam^fl/fl^* versus *Tfam^fl/flxMCM^* during functional resilience (2- and 20-week post-tamoxifen time points; Supplemental Figures 2–3), these transcripts were significantly downregulated at 32 weeks post-tamoxifen (Figure 6A). Further, transcript abundance of the helicase Twinkle (*Twnk*) and the mitochondrial polymerase gamma (*Polg*) significantly declined in *Tfam^fl/flxMCM^* hearts compared to *Tfam^fl/fl^* controls (Figure 6B). Taken together, prolonged loss of TFAM causes a loss of the core components of mtDNA transcription and replication machinery, and suppression of these systems likely contributes to the severe mitochondrial defects and cardiomyopathy (Figure 2) observed with long-term TFAM deletion.

**Figure 6.**
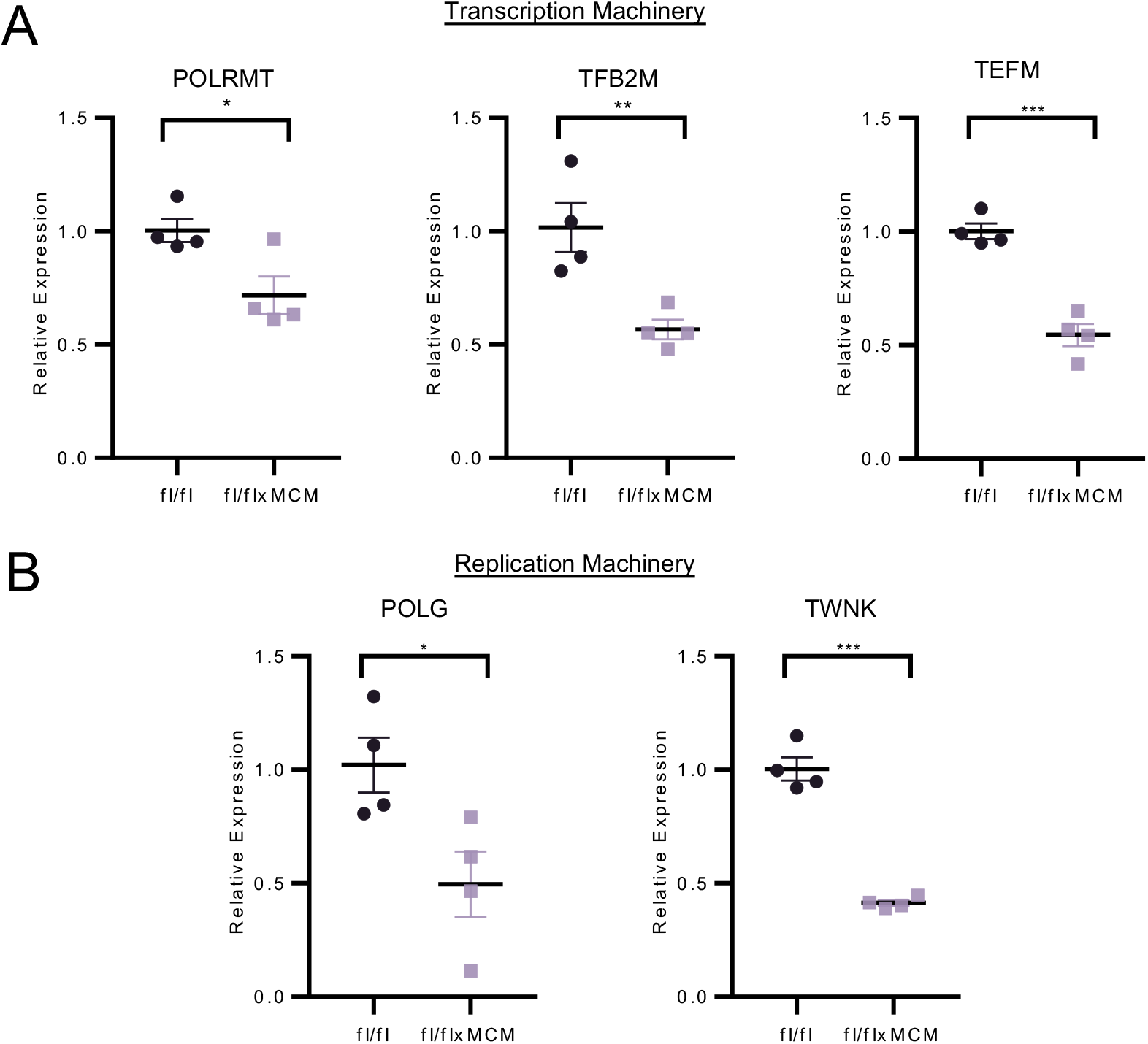
Downregulation of the mtDNA transcription and replication machinery with prolonged TFAM loss. (A) RT-PCR analyses of components of the mtDNA transcription machinery (*Polrmt, Tfb2m, Tefm*). (B) RT-PCR analyses of components of the mtDNA replication machinery *(Polg* and *Twnk*). Values presented as mean ± SEM. Student’s t test was used for statistical analysis. P<0.05 were considered significant. P<0.05 (*), P<0.01 (**), and P<0.001 (***).

## DISCUSSION

We conducted the first study of the functional consequences of mtDNA transcription and maintenance system defects in the adult heart. Our novel *Tfam^fl/flxMCM^* model of inducible TFAM deletion in adult mouse cardiomyocytes revealed surprising contrasts to phenotypes observed with systemic TFAM deletion (embryonic lethality) (21) or myocardium-specific TFAM deletion in embryonic or early postnatal life (cardiomyopathy, shortened lifespan) (23, 24). Indeed, inactivating *Tfam* in adult mouse cardiomyocytes resulted in an unexpected and extended period of cardiac functional resilience, wherein mitochondrial function, mtDNA content, and cardiac ventricular performance were preserved despite the loss of TFAM protein. While this functional resilience delayed disease onset, long-term deletion ultimately resulted in mitochondrial dysfunction and mitochondrial cardiomyopathy. These distinct phases of the cardiac response to TFAM inactivation illuminate unique features of adult heart mitochondria.

During functional resilience, while respiratory chain and cardiac performance were functionally indistinguishable between control and TFAM-null animals, key changes were induced by TFAM deletion. First, we observed derangements in mitochondrial organization in *Tfam^fl/flxMCM^* hearts at a time point when respiratory chain function was unchanged compared to *Tfam^fl/fl^* control mitochondria, suggesting that changes in mitochondrial morphology precede mitochondrial dysfunction. Interestingly, recent work with stimulated emission depletion nanoscopy of mitochondria revealed a highly ordered arrangement of mitochondrial nucleoids in relation to cristae, where nucleoids specifically occupy spaces separating groups of cristae (33). This finding suggests specific regulation underlying nucleoid-cristae positioning. In addition, within the nucleoid, TFAM interacts with mitofilin/Mic60, a critical component of the mitochondrial contact site and cristae organization system (MICOS) complex, which participates in cristae remodeling (34). Mitofilin deletion causes aberrant mitochondrial morphology with abnormal cristae structure and mtDNA depletion. While the mechanisms of mitofilin deletion-mediated mtDNA loss are not fully understood and may involve defects in nucleoid segregation during organelle division, our observations of abnormal mitochondrial morphology in the absence of both TFAM protein and overt mitochondrial energetics defects supports a model of TFAM-dependent regulation of mitochondrial morphology in the adult heart.

In addition to TFAM-dependent changes in mitochondrial ultrastructure, functional resilience to TFAM inactivation in adult cardiomyocytes was also marked by the surprising ability of *Tfam^fl/flxMCM^* mitochondria to maintain normal levels of mitochondrial respiration and a striking disconnect between TFAM-dependent mtDNA transcription and mtDNA content. Extensive literature implicates TFAM as a transcription factor for mtDNA and regulator of mtDNA content (as transcription generates the RNA primer to initiate mtDNA replication) and mtDNA compaction. In vitro and in vivo, TFAM loss induces coincident decreases in mtDNA transcription and copy number; in contrast, in vivo TFAM overexpression produces increased mtDNA copy number (19). Yet, our study revealed that TFAM protein levels do not automatically determine mitochondrial genome abundance in the adult heart: during functional resilience, *Tfam^fl/flxMCM^* hearts displayed preserved mtDNA content despite TFAM loss. Additionally, studies of mtDNA turnover rates in rodent heart calculated mtDNA half-lives ranging from 6.7 days (35) to 350 days (36). Our data support a model of a longer turnover time in adult cardiomyocytes (37), which may allow for mitochondrial genome abundance to be preserved following TFAM deletion.

Because mtDNA replication initiation requires transcription-based priming, our observations of unchanged mtDNA levels and preserved mitochondrial function in *Tfam^fl/flxMCM^* cardiac mitochondria initially led us to hypothesize that mitochondrial transcription would be similarly unaltered. However, decreases in mitochondrial transcript levels in *Tfam^fl/flxMCM^* hearts were an early response following induction of TFAM inactivation (decreases in transcript levels observed as early as 2 weeks post tamoxifen). Thus, acute deletion of TFAM in adult cardiomyocytes impairs mitochondrial transcription, and we postulate that mitochondrial translation—which was unchanged in *Tfam*-deleted versus control mitochondria during functional resilience—contributes to maintaining mitochondrial function. Ultimately, *Tfam^fl/flxMCM^* mitochondrial function might be subject to threshold effects, whereby declining mitochondrial transcript levels due to *Tfam* inactivation may not manifest as impaired mitochondrial translation until total transcripts decrease below a certain level. As our data suggests that the mitochondrial RNA degradosome is unaltered with TFAM deletion, our data may further support a model where transcript levels in cardiac mitochondria are in excess and mitochondrial translation serves as a response to sustain mitochondrial function in the face of acute defects to the mtDNA regulatory machinery (38).

How *Tfam^fl/flxMCM^* hearts transition from functional resilience to cardiomyopathy is perplexing, and complete delineation of this signaling pathway is beyond the scope of the present study. However, we found that long-term cardiomyocyte TFAM inactivation (32-weeks post-tamoxifen) downregulated the expression of the core mitochondrial transcription and mtDNA replication machinery. Interestingly, expression of these factors was unaltered during functional resilience (Supplemental Figure 2), suggesting that TFAM deletion-induced downregulation of the mitochondrial transcription and replication machinery is a specific feature of prolonged cardiomyocyte TFAM loss. Moreover, impaired mtDNA transcription and replication likely potentiates the decline in mitochondrial transcripts and genome abundance and may serve as a feed-forward mechanism underlying the mitochondrial and cardiac dysfunction observed with prolonged TFAM ablation.

In summary, we found that loss of TFAM elicits different effects on mtDNA transcription, replication, and maintenance systems in adult versus embryonic/postnatal hearts. While initiating *Tfam* deletion at either life stage causes cardiomyopathy, the differences in timing of disease onset and development are striking. A possible explanation for these differences—early and rapid onset of cardiomyopathy and shortened lifespan in embryonic/postnatal mice, versus functional resilience and slow disease onset in adult—is the inherent differences between these two stages: the embryonic/postnatal heart is a developing tissue undergoing active growth and proliferation, while the adult heart is terminally differentiated. Metabolic requirements, mitochondrial function, and even mitochondrial morphology differ between these two states—with the more fragmented mitochondria of the developing cardiomyocytes maturing into a highly interconnected reticulum of the differentiated cardiomyocytes, (28, 39, 40). Moreover, TFAM may have a critical role in cardiomyocyte proliferation (22). Thus, it is possible that cells undergoing active growth and proliferation have a greater dependence on TFAM function and could require higher rates of TFAM-facilitated mtDNA transcription and synthesis than terminally-differentiated cardiomyocytes.

Taken together, our study highlights the need to delineate mitochondrial maintenance pathways in both developing and differentiated tissues and organ systems, particularly because mitochondrial dysfunction is implicated in aging and disease.

## MATERIALS AND METHODS

### Animals

*Tfam^fl/fl^* loxP targeted mice were obtained from The Jackson Laboratories (stock #026123) and were generated as described in (21). *Tfam^fl/flxCre^* mice were generated by crossing *Tfam^fl/fl^* mice to animals expressing a Cre recombinase under the control of the alpha myosin heavy chain promoter (Cre) (29). The *Tfam^fl/flxMCM^* model of tamoxifen-regulated cardiac *Tfam* ablation was constructed by crossing *Tfam^fl/fl^* mice to animals expressing a tamoxifen inducible Cre recombinase under the control of the cardiomyocyte specific alpha-myosin heavy chain promoter (MCM) (30). *Tfam* deletion was induced in 8-week-old *Tfam^fl/flxMCM^* animals by intraperitoneal injections of tamoxifen (25 mg/kg for 5 consecutive days). Control animals, harboring either the *Tfam* targeted locus (*Tfam^fl/fl^*) or MCM transgene alone, were subjected to the same tamoxifen dosing regimen. All animal experiments were approved and performed in accordance to Emory University’s IACUC.

### Echocardiography

Mice were anesthetized with isoflurane (1.5%) with body temperature maintained at 37°C and measurements were performed using a Vevo 2100 Imaging System (Visual Sonics) equipped with a MS-400 transducer. M-mode measurements were taken of the parasternal short axis view. Systolic and diastolic ventricular chamber dimensions, ventricular wall and septal thicknesses were assessed, and fractional shortening was calculated using the VevoLab software.

### Electrophoresis and immunoblotting

For western blots, total protein extracts were prepared from hearts solubilized in radioimmunoprecipitation assay (RIPA) buffer supplemented with a combined protease and phosphatase inhibitor cocktail (Thermo Scientific). Mitochondrial protein extracts were prepared by solubilizing isolated cardiac mitochondria in the same RIPA buffer. For Western blotting of adult cardiomyocytes isolated from hearts by Langendorff perfusion with liberase blendzyme (0.25 mg/mL) and trypsin (0.14 mg/mL) in Krebs-Henseleit buffer as previously described (41). Proteins were reduced and denatured in Laemmli buffer, resolved on 10% SDS-PAGE gels, transferred to PVDF membranes, immunodetected with antibodies, and imaged using a ChemiDoc system (BioRad). Primary antibodies used in the study were: anti-TFAM (Abcam, ab131607, 1:1000), anti-Porin (Abcam, ab14734, 1:1000), and anti-GAPDH (Fitzgerald, 10R-G109A, 1:10000).

### mtDNA copy number analysis

Total DNA was prepared from snap-frozen heart tissue using the DNAeasy Kit (Qiagen). Quantification of relative mtDNA copy number was conducted by measuring the mtDNA/nDNA ratio using quantitative PCR. Primers for mouse mtDNA (Forward, CTAGAAACCCCGAAACCAAA; Reverse, CCAGCTATCACCAAGCTCGT) and the beta-2-microglobulin nuclear DNA (Forward, ATGGGAAGCCGAACATACTG; Reverse CAGTCTCAGTGGGGGTGAAT) were used as previously described (42). The qPCR assay was performed using the LightCycler 480 SYBR Green I Master (Roche) using Applied Biosystems QuantStudio 6 Flex Real-Time PCR System. Relative mtDNA copy number was calculated using the ΔΔC(t) method.

### RT-PCR

Total RNA was extracted from heart tissue using the RNeasy Fibrous Tissue Mini Kit (Qiagen) and cDNA was generated using the High Capacity cDNA Reverse Transcription Kit (Applied Biosystems). RT-PCR was performed on a QuantStudio 6 Flex Real-Time PCR System (Applied Biosystems) with LightCycler 480 SYBR Green I Master (Roche). Primer sequences were obtained from PrimerBank (43). ΔΔC(t) was used to quantify the fold change of the target genes. The primer sets used were: MTATP6 (Forward, CCTTCCACAAGGAACTCCAATTTCAC; Reverse, CTAGAGTAGCTCCTCCGATTAGGTG), MTCO1 (Forward, GCAGGAGCATCAGTAGACCTAAC; Reverse, GGAGTTTGATACTGTGTTATGGCTGG), MTCYTB (Forward, CTACTGTTCGCAGTCATAGCCAC; Reverse, CCAATATATGGGATGGCTGATAGGAG), MTND5 (Forward, GGCCTATTAATCGCAGCTACAGG; Reverse, GTAGTAGTGCTGAAACTGGTGTAGG) (25), MTND6 (Forward, ATGTTGGAAGGAGGGATTGGG; Reverse, TACCCGCAAACAAAGATCACC) (44), POLRMT PrimerBank ID 27369780a1 (Forward, GGCCCATCTTGCATTCTAGGG; Reverse, CAGGCAACGGCTCTATATTGAAG), TFB2M PrimerBank ID 6680223a1 (Forward, CCGCGTGCTGAGCATAAATC; Reverse, ACTGCACTAAGAGGTCCTGTG), TEFM PrimerBank ID 18044017a1 (Forward, CAACAAATGAGATGTGGCGATCA; Reverse, ACGAGCTTCTTACCAGCTATGA), POLG PrimerBank ID 8567392a1 (Forward, GAGCCTGCCTTACTTGGAGG; Reverse, GGCTGCACCAGGAATACCA), PNPT1 PrimerBank ID 12835817a1(Forward, AATCGGGCACTCAGCTATTTG; Reverse CAGGTCTACAGTCACCGCTC), SUPV3l1 PrimerBank ID 31088872a1 (Forward, GTGCAGCTCATGTGGACGATT; Reverse, GGGTGGTATCCTCAAGTCACTG) TWNK (Forward, GCCACGTGACTCTGGTCATTC; Reverse, CCATCAAAGCGATTCTTGGACA) (20), and RPS20 (Forward, AACAAGTCGGTCAGGAAGC; Reverse, TCCGCACAAACCTTCTCC).

### Transmission electron microscopy

Mouse heart samples were fixed in 2.5% glutaraldehyde in 0.1 M cacodylate buffer (pH 7.4), post-fixed in 1% osmium tetroxide, and embedded in epoxy resin. Ultrathin sections (80-90 nm) were cut with a Leica EM CU6 microtome and counterstained with uranyl acetate and lead citrate. Sections were imaged on a JEOL JEM-1400 transmission electron microscope (Tokyo, Japan) equipped with a Gatan US1000 CCD camera (Pleasanton, CA).

### Mitochondrial oxygen consumption

Cardiac mitochondria were prepared by differential centrifugation as previously described (45). Oxygen consumption was measured on isolated heart mitochondria using an Oxytherm System (Hansatech) as described (46). Briefly, mitochondria were suspended in respiration buffer containing 120 mM KCl, 5 mM MOPS, 0.1 mM EGTA, 5 mM KH_2_PO_4_, 0.2% BSA, 10 mM glutamate, and 2 mM malate. Basal ADP-stimulated respiration was initiated with the addition of 0.5 mM ADP, uncoupled respiration was induced with the addition of 5 μM FCCP, and non-mitochondrial oxygen consumption was assessed with the addition of KCN. Basal and maximal uncoupled mitochondrial oxygen consumption rates were obtained by subtracting KCN-non-mitochondrial respiration, followed by normalization to protein concentration.

### Histology

Hearts were fixed in 10% formalin and embedded in paraffin. Paraffin-embedded tissues were sectioned (7 μm) and stained with hematoxylin and eosin (H&E) and Masson’s Trichrome. Images were captured using a Nanozoomer 2.0-HT whole slide scanner (Hamamatsu). Myocyte cross-sectional area was quantified on H&E stained sections using NIH Image J software.

### In organello translation assay

In organello translation assays were performed with a modified protocol based on (47). Briefly, cardiac mitochondria were suspended in mitochondrial isolation buffer (25 mM sucrose, 75 mM sorbitol, 100 mM KCl, 1 mM MgCl_2_, 0.05 mM EDTA, 10 mM TrisHCl, and 10 mM K_2_HPO_4_, pH 7.4) supplemented with 10 mM glutamate, 2.5 mM malate, 1 mM ADP, 1 mg/mL fatty acid-free bovine serum albumin, and 100 μg/mL cycloheximide and incubated with 100 μM L-azidohomoalanine (AHA; Click Chemistry Tools) and 10 μM methionine-free amino acid mixture (Promega). The Click-iT reaction was performed with the Click-&-Go Protein Reaction Buffer Kit (Click Chemistry Tools) and AF488 alkyne (Click Chemistry Tools) according to the manufacturer’s instructions. Proteins were resolved by SDS-PAGE and imaged using a ChemiDoc MP Imaging System (BioRad).

### BN-PAGE

BN-PAGE followed by immunoblotting for respiratory complexes was performed as described in (48, 49). Briefly, isolated heart mitochondria were solubilized with digitonin (Sigma Aldrich, 6g/g digitonin/protein ratio) and resolved on a 4-12% Bis-Tris polyacrylamide gradient gel (Genscript). Proteins were transferred to PVDF membranes and immunodetection of respiratory chain complexes was performed using the Total OXPHOS Blue Native WB Antibody Cocktail (Abcam, ab110412, 1:250). The membrane was imaged using a Chemidoc XRS+ System (BioRad).

### Statistics

All results are presented as mean ± SEM. Statistical significance between 2 groups was determined by Student’s t-test, with P<0.05 considered significant.

### Study approval

All experiments involving mice were approved by the IACUC at Emory University, approval number PROTO201700174.

## Author Contributions

JQK and NG wrote the manuscript. NG, TM, JNP, and JQK performed experimentation. JQK, and NG analyzed the data. JQK designed the study and provided experimental oversight.

## Acknowledgements

Electron microscopy for this study was supported by the Robert P. Apkarian Integrated Electron Microscopy Core (RPAIEMC), which is subsidized by the Emory University School of Medicine and the Emory College of Arts and Sciences. Additional support for electron microscopy was provided by the Georgia Clinical & Translational Science Alliance of the National Institutes of Health under award number UL1TR000454. Echocardiography for this study was supported in part by the Animal Physiology Core, which is subsidized by Emory University and Children’s Healthcare of Atlanta. Additional support was provided by the Office of the Director of the National Institutes of Health under Award Number S10OD021748. Histology for this study was supported in part by the by the Cancer Tissue and Pathology shared resource of Winship Cancer Institute of Emory University and NIH/NCI under award number P30CA138292.

---

**Supplementary Figure 1.**
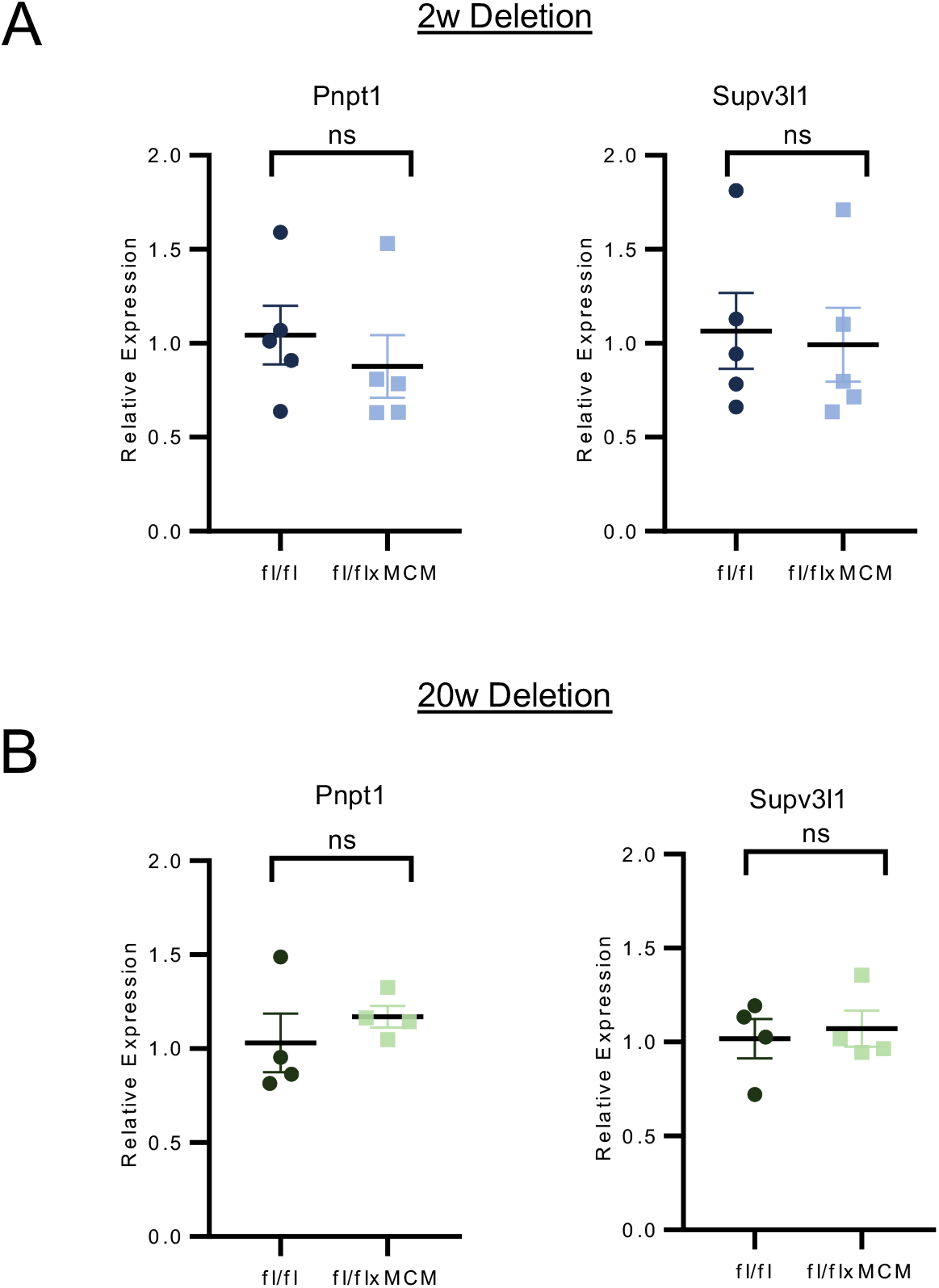
Expression of the mitochondrial degradosome is unaltered with cardiac TFAM deletion during functional resilience. RT-PCR of cardiac *Pnpt1* and *Supv3l1* expression assessed at (A) 2 weeks and (B) 20 weeks following tamoxifen administration.

**Supplementary Figure 2.**
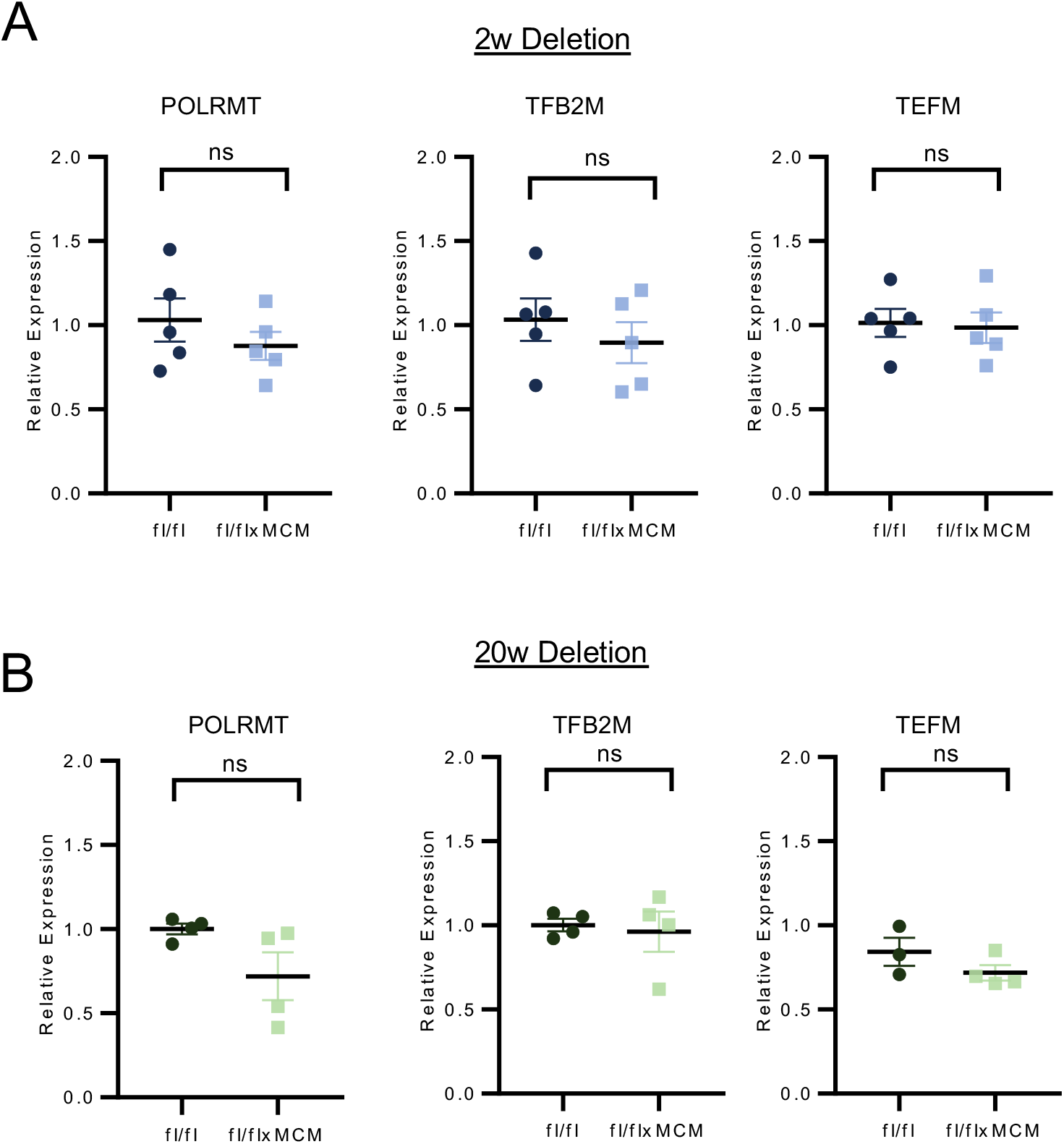
Expression of core components of the mitochondrial transcriptional machinery is unaltered during functional resilience to TFAM deletion. RT-PCR of cardiac *Polrmt, Tfb2m*, and *Tefm* transcript expression assessed at (A) 2 weeks and (B) 20 weeks following tamoxifen administration.

**Supplementary Figure 3.**
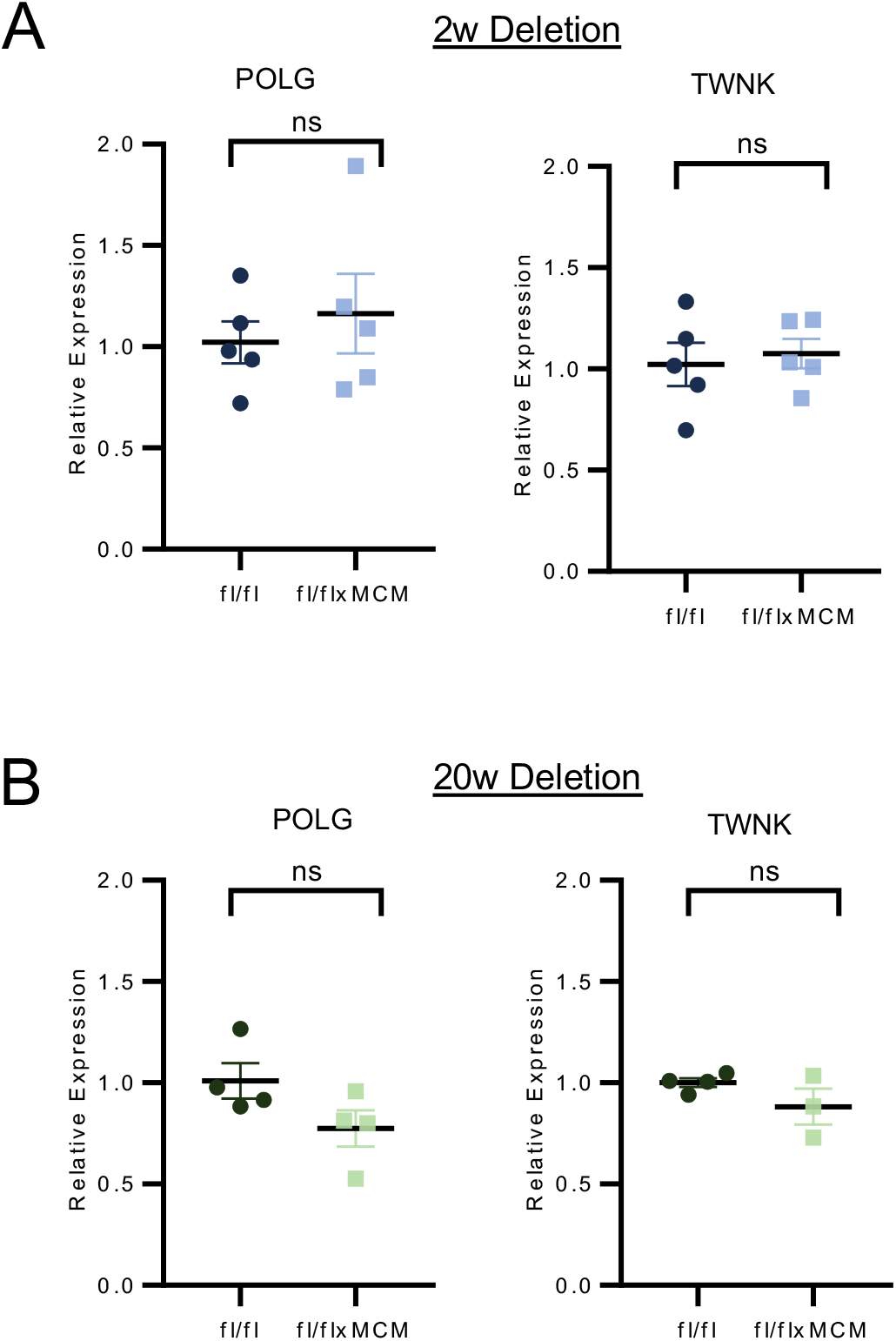
Expression of core components of mtDNA replication is unchanged during functional resilience to TFAM deletion. RT-PCR of cardiac *Polg*, and *Twnk* transcript expression assessed at (A) 2 weeks and (B) 20 weeks following tamoxifen administration.

## Notes

### Competing Interest Statement

The authors have declared no competing interest.

